# On the proportional abundance of species: Integrating population genetics and community ecology

**DOI:** 10.1101/223529

**Authors:** Pablo A. Marquet, Guillermo Espinoza, Sebastian R. Abades, Angela Ganz, Rolando Rebolledo

## Abstract

The frequency of genes in interconnected populations and of species in interconnected communities are affected by similar processes, such as birth, death and immigration. The equilibrium distribution of gene frequencies in structured populations is known since the 1930s, under Wright’s metapopulation model known as the island model. The equivalent distribution for the species frequency (i.e. the species proportional abundance distribution (SPAD), at the metacommunity level, however, is unknown. In this contribution, we develop a stochastic model to analytically account for this distribution (SPAD). We show that the same as for genes SPAD follows a beta distribution, which provides a good description of empirical data and applies across a continuum of scales. This stochastic model, based upon a diffusion approximation, provides an alternative to neutral models for the species abundance distribution (SAD), which focus on number of individuals instead of proportions, and demonstrate that the relative frequency of genes in local populations and of species within communities follow the same probability law. We hope our contribution will help stimulate the mathematical and conceptual integration of theories in genetics and ecology.

## Introduction

Ever since the evolutionary synthesis, population genetics theory has been integrated, to different extents, into different disciplines within biology including systematics and ecology. This later integration took off with the development of theoretical formulations relating the processes that drive changes in numbers of individuals within age-structured populations, with changes in the fitness of different genotypes^1,2^. Yet further integration was achieved with the emergence of the new ecological genetics spoused by Antonovics^3^, one of whose tenets was that “*Forces maintaining species diversity and genetic diversity are similar. An understanding of community structure will come from considering how these kind of diversity interact*.” More recently, the emergence of community genetics^4^ has reinvigorated the search for connections between population genetics and community ecology, along with the realization that there is a striking similarity between processes driving changes in the abundance and diversity of species within communities and genes within populations^5,6^.

The recent development of neutral approaches to the study of ecological systems^7–10^ have provided a renewed emphasis upon the value of theory and stochasticity in ecology^11–14^ and a locus for the further integration of genetical and ecological theories^15,16^. By merging the mathematical and statistical tools developed by population geneticists with the neutrality approach, neutral theory in ecology allows us to better understand the factors affecting the abundance and distribution of species^15–19^. But there is a major barrier to this integration, while population geneticists pioneered the use of diffusion approximations (i.e. a continuous process) to the understanding of processes affecting gene frequencies^20^, ecologists have favored to work with the distribution of the number of individuals across species (i.e. a discrete process) or SAD^8,21–23^ (but see^24^). It is not surprising then that the answer for the abundance of species within communities (i.e. Fisher’s Log-series,^8^) is different from that for gene frequencies within populations (i.e., a Beta distribution^25,26^). In this contribution, we aim at filling this gap in knowledge by analyzing the distribution of species abundances as a continuous process (i.e. using a diffusion approach). To do so we focus on the proportional abundance of species instead of the number of individuals. We show that if one assumes that birth and death rates follow the functional form used in neutral theory citemckane2000,volkov2003 the stationary distribution for the species proportional abundance distribution (SPAD), the same as for genes, is a beta distribution with parameters *α* and *β* that quantify the relative importance of immigration and speciation, respectively, in relation to stochastic fluctuations. We show that this distribution provides a good description of empirical data and applies across a continuum of scales.

## The model

We model the community as an open system, and as such we do not distinguish two spatial scales in our system, as usually done in neutral models, as the one proposed by Volkov et al.^8^, but a continuum of scales, which are defined by the observer of the system when studying it. The system could be, for example, a 50 ha plot in a tropical forest or a 1m^2^ plot in the intertidal. What is important is to realize that once the observer defines the spatial scale of the system, it defines a boundary or an inside and an outside, where the focal system is embedded (Figure 1). We call this observer defined scale the focal community that is embedded into a bath or environment with whcih it interacts. The focal community dynamics is driven by birth and death processes and by immigration from the outside. We do not explicitly consider speciation as this is subsumed into the immigration process^11^. Indeed the spatial scale of analysis is to some extent dictated by which is the dominant process adding new species to a given focal community; immigrations of individuals from species not yet found in the focal community but somewhere else in the bath, or new species arising through speciation within the focal community. If the later is the dominant process, then the spatial scale is likely to be large, since all species in the potential pool are already present and the only way a new species can arrive would be through speciation. Similarly, the processes that remove individuals and species from the focal community include death and emigration towards the bath or environment. To model the dynamics of this focal community we used the diffusion approximation of birth and death processes independent of a focal community size *J*. By community size we mean the total number of individuals regardless of species identity.

**Figure 1.**
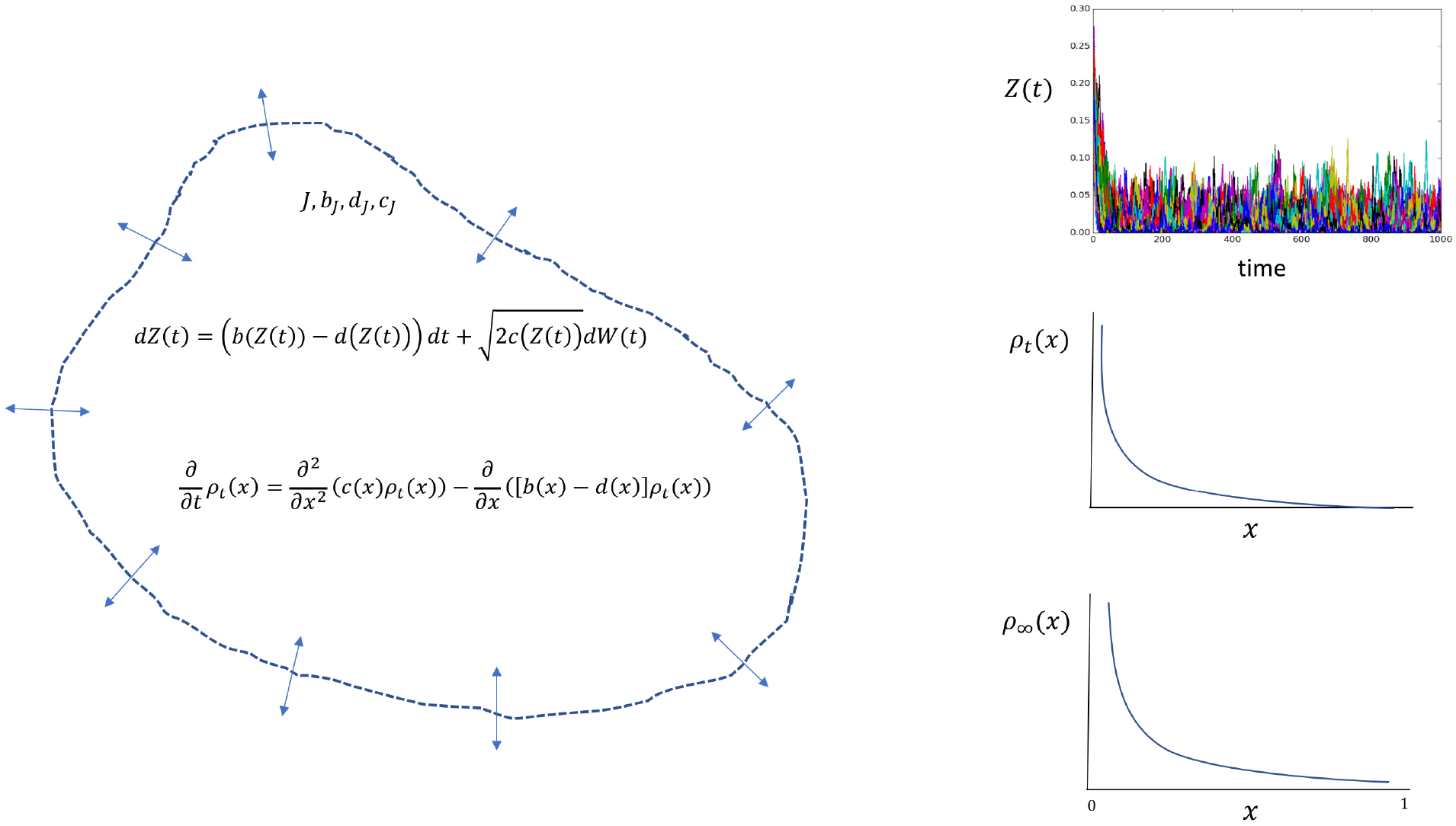
Diagrammatic description of the diffusion approach taken in this contribution. This approach assumes the existence of a focal community (the white area delimited by a discontinuous line) of size *J*, and where *N_J_*(*t*) denotes the number of individual of a given species within it. The abundance of any species in this focal community follows a birth death process, with rates *b_J_*, *d_J_* and *c_J_*. However, since we are interested in the proportion of individuals instead on their numbers, we introduce the process *Z*(*t*) or stochastic proportional abundance. It is shown that as *J* → ∞, *Z*(*t*) converges to a diffusion that satisfies the stochastic differential equation for *dZ*(*t*) with rates *b*(*x*), *d*(*x*) and *c*(*x*) (see Eq. (7–9)). At any given time the probability density of *Z*(*t*) is given by the Fokker-Planck equation associated to *ρ_t_*(*x*) (Eq.(10)). Further, when *t* → ∞ this probability density becomes stationary or invariant and is called ρ_TC_(x). We show that when *b*(*x*), *d*(*x*) and *c*(*x*) have a particular functional form (see Eq. (12–14)) the invariant distribution is a beta distribution (Eq. (15)). The Panels on the right show the simulation of trajectories for the diffusion process *Z*(*t*), the associated density at a given time *ρ_t_*(*x*) and the invariant distribution *ρ*_∞_(*x*).

Let *N_J_*(*t*) denote the number of living individuals of a given species within a focal community of size *J*, at time *t* ≥ 0 (so that *N_J_*(*t*) is less or equal to *J* for all *t*). This is assumed to be a birth and death process, with transition matrix *P*(*t*) = (*P_n,m_*(*t*); *n*,*m* = 0, …, *J*) (*n* and *m* denotes the number of individuals). For a small time increment *h*, this matrix satisfies as *h* → 0 for *n* ≥ 0

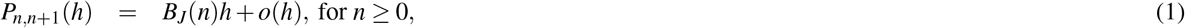

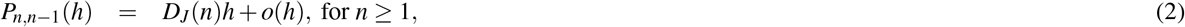

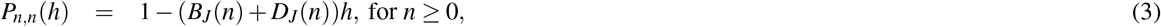

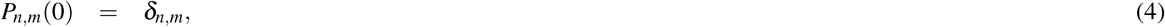

where *B_J_*(*n*) and *D_J_*(*n*) are the birth and death rates, respectively, 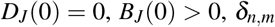, is the customary Kronecker delta, and *o*(*h*) denotes the Landau-symbol, which satisfies 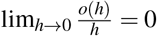. Here, in addition, we assume that these rates are decomposed as follows

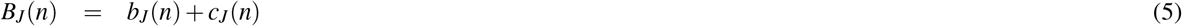

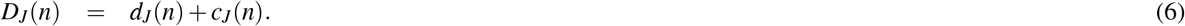

The terms *b_J_* and *d_J_* represent birth and death rates in the focal community, respectively, which will be asymptotically independent of *J*, while *c_J_* takes into account the variations on the above rates due to the interaction between the focal system and the environment wherein it is embedded, proper to an open system approach. Since we are interested in proportions *n/J*, we introduce the variable *x* = *n/J*, which takes values in {0, 1/*J*, 2/*J*, …, 1}, and analyze the behavior of the system as the size of the population grows indefinitely: *J* → ∞. At this stage it is important to state meaningful hypotheses for the previous rates for large *J*, as all changes of scales in the dynamics of the open system are driven by this community size.

We first assume that *b_j_* and *d_J_* will lead, respectively, to the *J*-invariant (or endogenous) birth and death rates of the focal system, that satisfy

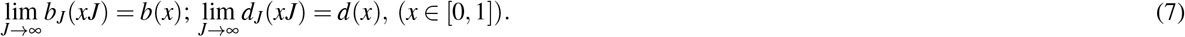

On the contrary, the rate *c_J_*, should vary significantly with *J*, however, we require that it satisfies

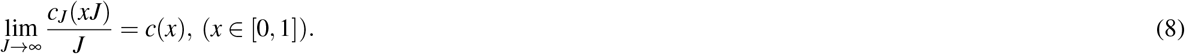

We can now define the stochastic process 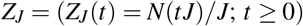 that we call the stochastic proportional abundance. This family of processes has a limit 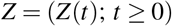 as *J* → ∞, that corresponds to a diffusion process satisfying the stochastic differential equation (see Supplementary Information)

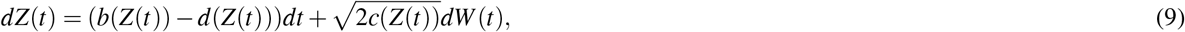

where *W*(*t*) denotes a Brownian motion.

It is worth noticing (see Supplementary Information also) that the process 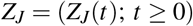 converges in distribution towards a diffusion process 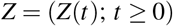 as proven in^27^, and so, any continuous functional *F*(*Z_J_*) of the trajectory of *Z_J_* converges in distribution to *F*(*Z*). In particular, it is proved (see Supplementary Information) that for any values 0 < *a* < *b* ≤ 1, it holds

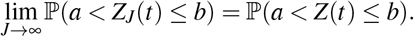

where 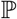 is the probability defined on the set of all trajectories of the process.

Correspondingly, the Fokker-Planck equation associated with the probability density *ρ_t_*(*x*) of *Z*(*t*), is given by

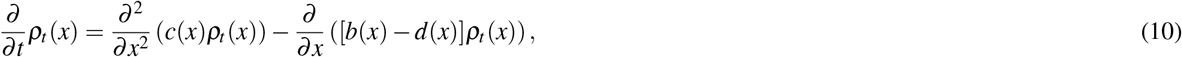

with the additional condition that 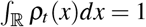. The stationary solution ρ_∞_ is determined as the solution to the equation

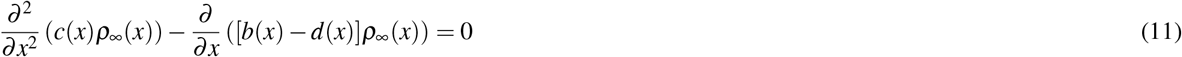

In order to find the stationary distribution we need to make a hypothesis for each of the rates *b*(*x*), *d*(*x*) and *c*(*x*), the simplest ones are that

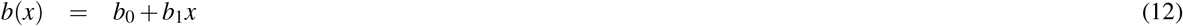

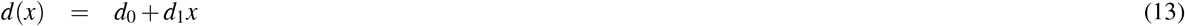

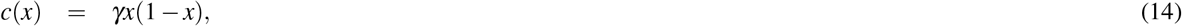

where *b_i_*, *d_i_*, (*i* = 0,1), and *γ* are positive constants. Under these hypotheses (see Supplementary Information) the stationary solution takes the form of a typical Beta distribution

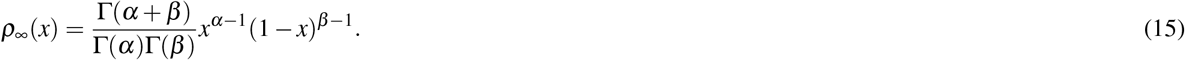

Then an elementary computation shows that (15) provides a solution to (11) with

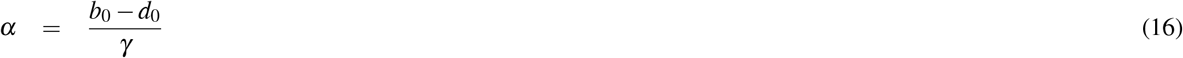

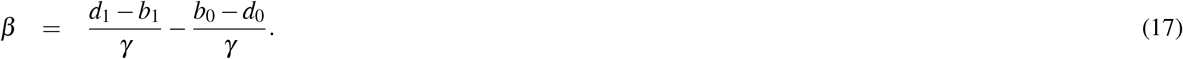

In Figure 1, we provide a diagrammatic version of the main steps taken in our derivation of the stationary Beta distribution. As an important particular case, let us use the rates proposed by McKane^28^ and used in the neutral theory model proposed by Volkov^8^, which in our framework, this will correspond to the following rates

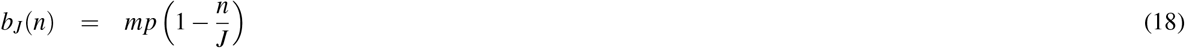

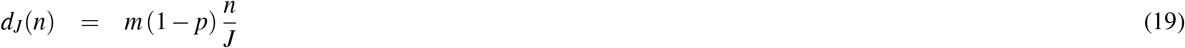

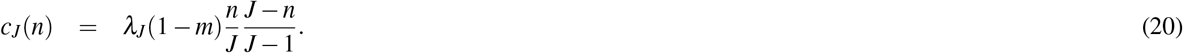

where *p* is the probability with which we choose individuals of a given species, and *m* denote a migration probability. In addition, we introduce the parameter *λ_J_* to keep track of fluctuations in demographic rates due to interactions between the focal system and the environment, for instance, as a consequence of temperature variations or due to other unknown biotic or abiotic variables. We assume that *λ_J_*/*J* → *λ* as *J* → ∞. Thus, letting *J* → ∞, one obtains the convergence towards the corresponding limits

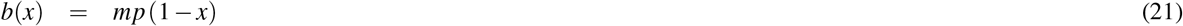

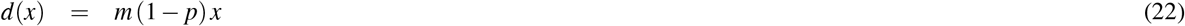

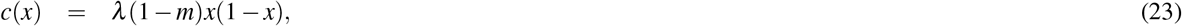

where *x* ∈ [0,1] (that is, *b*_0_ = *mp*, *b*_1_ = −*mp*, *d*_0_ = 0, *d*_1_ = *m*(1 − *p*), *γ* = *λ* (1 − *m*)).

Thus, under the above choice of coefficients, (9) becomes

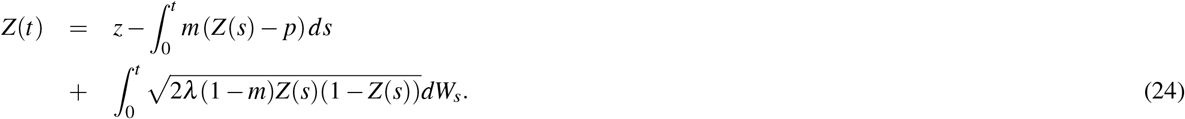

And *ρ*_∞_ has the form (15) with

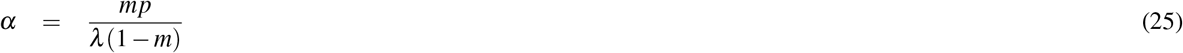

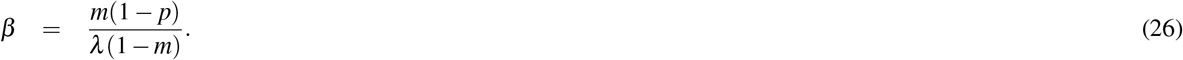

We interpret *α*, as quantifying the relative contribution of immigration of known species to the abundance of species already present in a focal community, while *β* quantifies the relative contribution of immigration of species not yet known in the focal community, that is, speciation. Notice that, both *α* and *β* are expressed in relation to the magnitude of the fluctuations induced by the interaction with the environment (i.e. *λ* (1 − *m*)).

When the probability with which we choose individuals of a given species is *p* = 1/*S*, where *S* denote the total number of species, *β* = *α* (*S* − 1) and thus (15) becomes

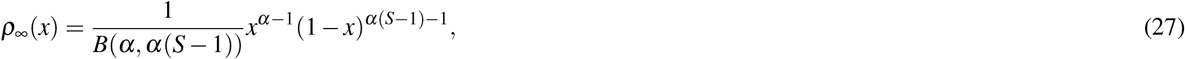

where 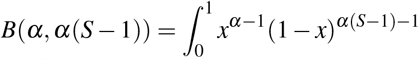 (normalization constant) and 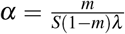 (see derivation of the Beta distribution in the Supplementary Information).

In Figure 2 we show the fit of (27) to several datasets including the Malayan butterflies and the Rothamsted Lepidoptera data originally used by Fisher^29^, tropical birds in Manu Park (Perú)^30^, tropical forests^31^, Fynbos shrublands^32^ and coral reefs^33^. In all cases the correlations between observed and fitted frequencies (expressed as proportional abundance) was highly significant (Table 1).

**Figure 2.**
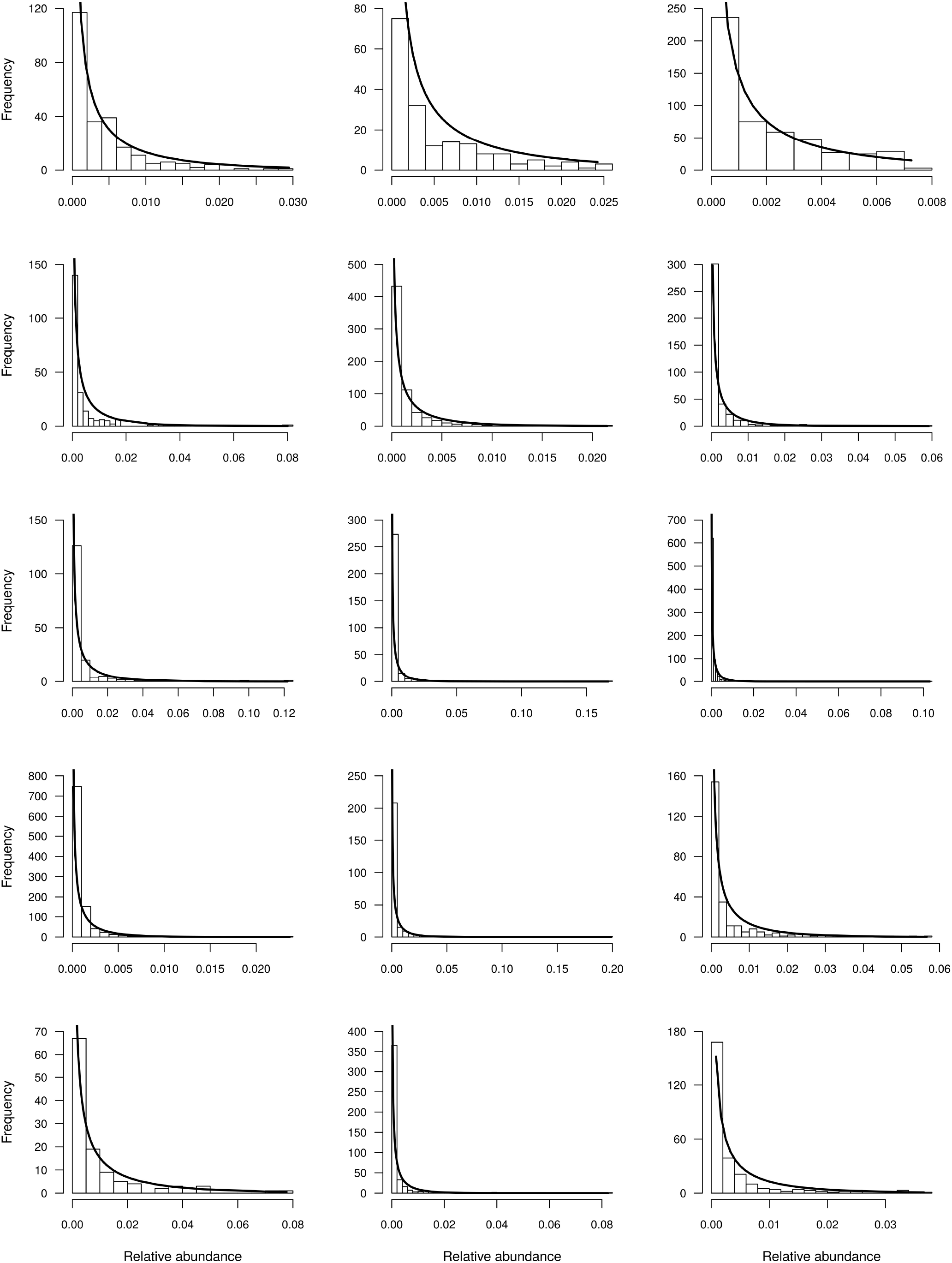
Fit of the Beta distribution to different animal and plant communities. From left to right, first row, Amazon birds, Lepidoptera, butterflies, second row Tropical trees and Coral reefs (communities 10, 12, 11, 6, 2 and 14 in Table 1 respectively. Third row Tropical trees. Fourth row Tropical trees, and Fynbos shrublands. Fifth row Fynbos shrubland and coral reefs (communities 1, 3, 4, 5, 7, 8, 9, 13 and 15 in Table 1 respectively)

**Table 1.** Fit of the Beta distribution (Eq. (27)) to fifteen plant and animal communities. Data for communities 1-6 comes from^31^, 7-9 from^32^ 10 from^30^, 11-12 from^29^ and 13-15 from^33^. The estimation of *α* and *β* was done by optimization based on the Nelder-Mead method implemented in the maximum likelihood function mle2, included in library bbmle for R. Comparison between observed and predicted frequency distribution were done using Pearson’ s correlationdone by optimization based on the Nelder-Mead method implemented in the maximum likelihood function mle2, included in library bbmle for R. Comparison between observed and predicted frequency distribution were done using Pearson’s correlation

| Community | S | J | $\alpha$ | $\beta$ | Pearson's r |
| --- | --- | --- | --- | --- | --- |
| 1 Sinharaja | 167 | 16936 | 0.2498 | 41.4668 | 0.915 |
| 2 Pasoh | 678 | 26554 | 0.3868 | 261.8370 | 0.978 |
| 3 Korup | 308 | 24591 | 0.2783 | 85.4514 | 0.945 |
| 4 Yasuni | 821 | 17546 | 0.4872 | 399.4604 | 0.967 |
| 5 Lambir | 1004 | 33175 | 0.4291 | 430.3599 | 0.987 |
| 6 Barro Colorado Island | 225 | 21457 | 0.2773 | 62.1201 | 0.897 |
| 7 Hangklip | 247 | 23756 | 0.2538 | 62.4323 | 0.927 |
| 8 Cederberg | 247 | 11561 | 0.3025 | 74.4140 | 0.849 |
| 9 Zuurberg | 114 | 8806 | 0.3709 | 41.9143 | 0.415 |
| 10 Terborgh | 245 | 1663 | 0.8796 | 214.6275 | 0.854 |
| 11 Fisher Butterflies | 501 | 3306 | 0.9877 | 493.8308 | 0.891 |
| 12 Fisher Lepidoptera | 180 | 2020 | 0.6976 | 124.8712 | 0.905 |
| 13 Dornelas Indo Pacific | 450 | 3779 | 0.6427 | 288.5661 | 0.840 |
| 14 Dornelas Papua New Guinea | 403 | 2520 | 0.8557 | 344.0007 | 0.864 |
| 15 Dornelas Solomon Islands | 268 | 1201 | 1.1268 | 300.8603 | 0.834 |

In Figure 3 we show the relationship between *α* and *β*. As expected, both are positively correlated, but more interestingly it is apparent that birds, butterflies and marine communities are characterized by large *α*, a measure of the importance of migration, as expected for open and highly connected systems where immigration in the form of dispersal could be the dominant processes accounting for the appearance of new individual each generation. Similarly, the Fynbos shrub dominated communities (7-9 in Table 1) are characterized by low *β*, which may be associated to low rates of speciation (but see^32,34^). Indeed, *β* is correlated to the biodiversity number θ of classical neutral theory (Pearson’s r=0.97,n=6, P<0.01, see inset in Figure 3), which is a function of speciation rate^7,8^.

**Figure 3.**
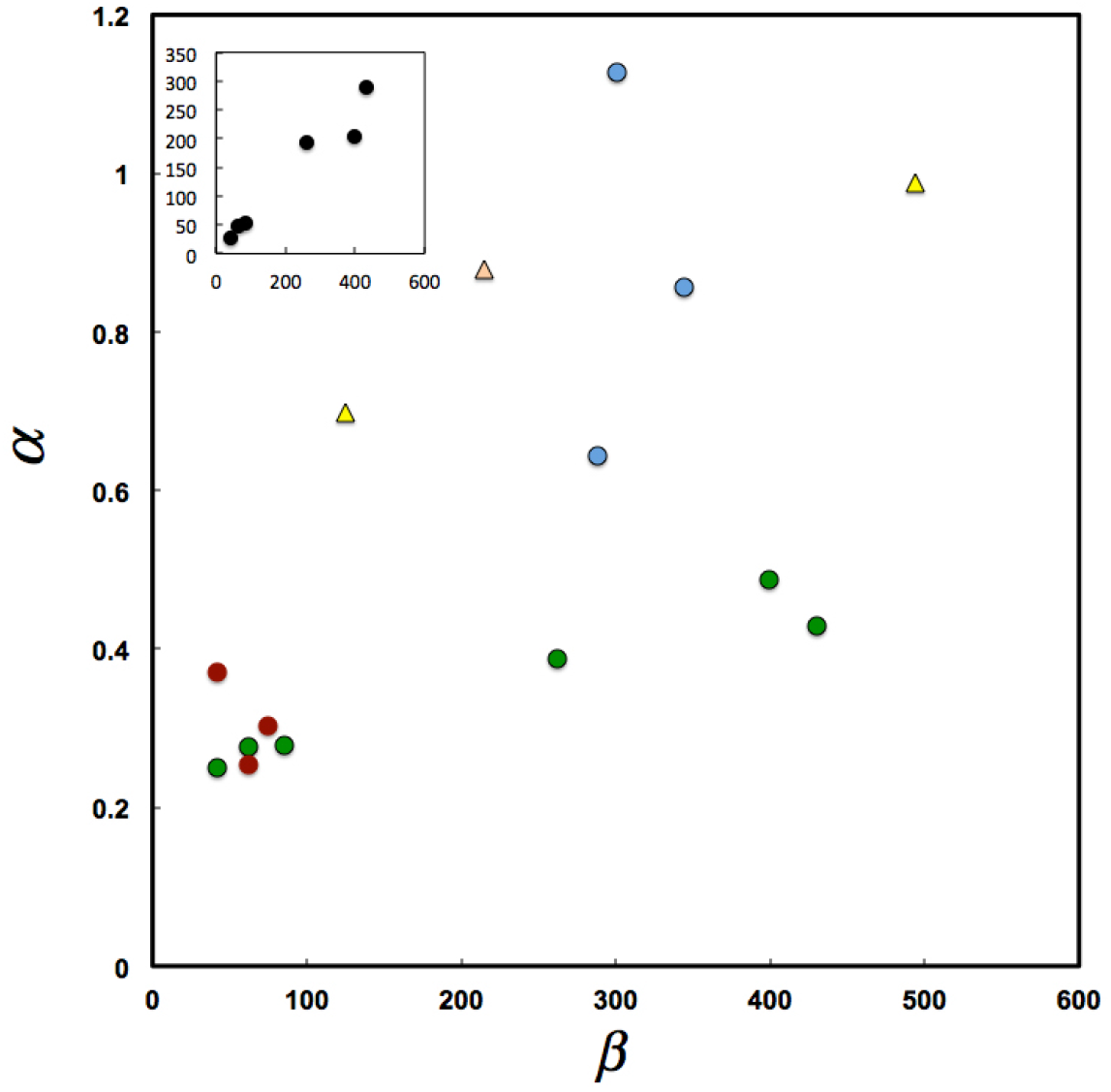
Relationship between parameters *α* and *β* for the communities shown in Table 1 (in blue Marine, in green Tropical Forest, in red shrublands, in light yellow butterflies and in strong yellow bird communities). In the inset the relationship between *β* (y axis) for forest communities 1-6 in Table 1, and the θ (x axis) parameter estimated in^31^ for the same forest communities.

Finally, in Figure 4a, we show simulations of the stochastic proportional abundance of species or trajectories of *Z*(*t*) in (24). Figure 4b is the plot of the confidence intervals around the mean 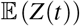, notice that the process rapidly converges to the long term average value. As we mentioned before, the density distribution *ρ_t_* of the stochastic proportional abundance, which corresponds to a neutral abundance at the rescaled time *t*, tends to a stationary distribution *ρ_∞_* as *t* → ∞. We can estimate *ρ*_∞_ by sampling the trajectories of *Z*(*t*) after a large number of generations (e.g. *t* = 1000) represented by the histogram in Figure 4c, which is in good agreement with the limit beta distribution density *ρ*_∞_.

**Figure 4.**
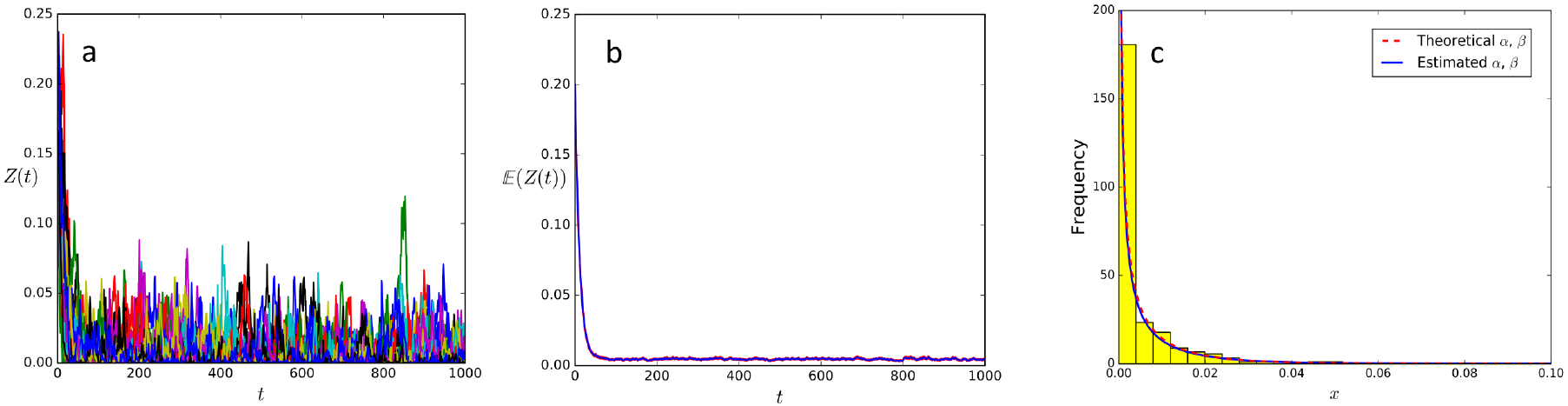
a) Simulation of 225 trajectories (only 50 are shown) using Eq.(24) with *λ* = 0.001585, *p* = 0.0044, *m* = 0.09 and an initial proportional abundance Z(0) equal to 0.2. b) Mean value of the observed trajectories and 95% confidence intervals. c) Histogram of the trajectories *Z*(*t*) for *t* = 1000 in (a), estimated Beta distribution (27) (continuous blue line) and the theoretical density *ρ*_∞_ (27) (red dashed line).

## Discussion

A key component of the evolutionary synthesis was the mathematical formalization of the processes driving changes in gene frequencies within Mendelian populations. Wright’s island model^25,35^ demonstrated that the frequency of neutral alleles in a local open population affected by mutation, migration and drift, will converge to a Beta steady-state distribution of allele frequencies. In light of our results, the equilibrium distribution of gene frequencies in a local population is equivalent to the frequency of different species in a local community or the Species Proportional Abundance Distribution (SPAD). Although this equivalence was expected, as both genes and species are affected by similar stochastic processes, it is a novel result since the equilibrium distribution of the SPAD was unknown, and previous results have either relied upon additional assumptions, such as density dependence^36^ or on approximations to the continuous limit^37,38^. Our results complement the efforts to understand the distribution of species abundances that have focused on changes in the numbers of individuals in different species (e.g.^7,8,29^) instead of the proportional abundance of species within communities. As far as we know, however, ours is the only continuous, neutral, and exact mathematical formulation derived from first principles. That is, based upon a birth death processes on the appropriately rescaled relative abundance process that, in the limit as *J* → ∞, is shown to satisfy the stochastic differential equation (9) in agreement with Rebolledo’s central limit theorems^27^ (see also Supplementary Information).

The general model for SPAD that we propose is based on a diffusion approach, as it has been used in population genetics to study the distribution of gene frequencies. Indeed Kolmogorov^39^, showed that the steady state distribution for allele frequencies (i.e. a Beta distribution) in Wright’s island model was the stationary distribution of the diffusion approximation. In this vein, we show that the stationary distribution for the species proportional abundance is a Beta distribution, but only if birth and death rates are of the form (12), which accommodates, as a particular case, the ones traditionally used in neutral models^8,28,31^.

Since the gamma distribution is the invariant distribution of a single species population following stochastic logistic growth^40,41^, it has been suggested as the most appropriate to describe SADs^42^. Interestingly, Fisher’s logarithmic series model is a Gamma type distribution. It is derived from Poisson sampling a population of S species (i.e. when the number of individuals sampled from any species is Poisson distributed) whose abundance follows a gamma distribution with shape parameter *k* = 0. As shown by Kempton^43^ if the sampled population consists of independent subpopulations each following a generalization of the Gamma model (i.e. *k* ≠ 0) then the resulting distribution will be a Beta distribution, as it is well know in statistics, and the resulting sampling distribution would be the generalized log series. Similarly, Engen and Lande^42^ show that under a stochastic logistic model with positive mean growth rate, the relative abundances of species would be Dirichlet distributed, which is the multivariate version of the beta distribution. Thus, the beta distribution has been around for a long time in ecology, here we show it is the invariant distribution associated to a diffusion process representing an open dynamical system under neutrality.

It is important to realize that the stochastic process described by *Z*(*t*) is of the Markov type since future changes depend on the present state, but not on the past history which led to this present state. Although this is a common assumption in ecological and evolutionary models, a large body of experimental data and analyses shows the importance of history (or memory) in affecting current states at the level of individuals, populations and lineages^44–46^. In this context it will be desirable to develop non-markovian models for neutral macroecology; after all, life is a historical process and the explicit consideration of history may be the simplest way of breaking the symmetry of neutrality.

If the variable *Z*(*t*) were discontinuous (i.e. if it were a measure of number of individuals instead of proportions) it will change in jumps due to birth, death, immigration and speciation processes and in this case the probability of a change during a small time interval (*t*, *t* + *h*) is small (of the order of magnitude *h*), but if a change occurs, it is of finite magnitude. In the diffusion approximation, during any time interval, however small, *Z*(*t*) undergoes some change, such that the probability that *Z*(*t* + *h*) − *Z*(*t*) > *ε* is of smaller order of magnitude than h. Continuity in this case, is possible only for large J as the number of event per time interval become continuous in rescaled time (i.e. *tJ*).

In genetics, where diffusion methods where first applied in the context of biology, the diffusion approximation was used to derive the distribution of allele frequencies under the process of migration, mutation, selection and drift (by themselves and in combination)^47^. Interestingly, in this area of inquiry, diffusion methods provided good approximations to model the evolution of finite populations^48^, even though its derivations requires *J* → ∞. In our case, the derivation of the beta distribution is based on two limits one for the number of individuals, and secondly, one in time. The order in which these limits are taken cannot be changed. Once the diffusion limit is obtained via *J* → ∞, the beta distribution is indeed obtained as a consequence of *t* → ∞. Since what we are analyzing is the evolution of individual abundance, a process that started with the origin of life, it is correct to assume that we are at the large *t* limit (even if we consider the time since the last major extinction event 66 million years ago) and thus the finding of a beta distribution should be common. In our case, the fits to finite focal communities seems remarkable, however we do not know how *J* affects the fit to our stationary solution and if there is a minimum *J* below which our approximation would seem inadequate. The issue get even more complex since the Beta distribution does not have a close form Maximum Likelihood estimator, which hinders the usability of the model in terms of estimating parameters of the distribution given the data, and testing hypotheses about them. An alternative solution is to use an approximation to the maximum likelihood, several of which are implemented in available packages such as R, Matlab and Scipy, and which provide accurate estimations of parameters (less that 3 percent bias) with sample size above 100^49^, or to estimate the coefficients of the diffusion process itself using the methods suggested by^50^ and simulate the stochastic process (9) to obtain the expected form of *ρ_t_* as shown in Figure (3) and then compare it to empirical ones. Although in strict terms *Z*(*t*) and its invariant apply to one species, the neutrality assumption allow us to use *ρ*_∞_ as a good hypothesis for multispecies assemblages. In this context we show in the Supporting Information (Figs. S1-3) that the parameters of the Beta distribution *β*, *α* can be estimated with little error when simulating 200 trajectories of *Z*(*t*) (see also Figure 4c), which as a first approximation we consider as a proxy for 200 species under neutrality. Finally, if the steady state assumption in (11) does not hold, due to perturbations or in the case of a newly colonized habitat, then we will be observing *ρ_t_* and its functional form can be explored through simulations (codes provided upon request). These are important issues that require further investigation to increase the applicability of the diffusion approximation herein provided.

Our diffusion approximation is based upon the paradigm of open dynamical systems, whereby we try to understand the behavior of a focal system, or focal community, in the context of an environment or bath with which it interacts; an approach that has been mostly developed for open quantum systems^51^. Since we are only able to specify the dynamics of our focal system, which is th one we study and develop theories an hypothesis about, everything we do not know about it is specified in the fluctuations represented by the noise term in the stochastic differential equation (9), whose intensity is dependent upon the the value of *c*(*x*). In this respect, our model can accommodate both neutral and non-neutral processes, with the latter being included in the noise term. In the particular case we explored, using transition rates as in^28^, the core of the dynamics is neutral at the level of the focal system but everything else that could potentially impact upon the dynamics of the local systems, either neutral or not, will be capture in the fluctuations induced by the interaction with the reservoir and included in the Brownian noise term. It is important to notice that we assume that these fluctuations act at comparable time scales, if this were not the case (as it is likely since immigration is faster than speciation) the addition of a different time scale in the form of fluctuations following a Poisson distribution may be in order. In this case we would arrive to a *Lévy* type diffusion process.

One of the problems of our derivation is that there are no comparable models against which to contrast its performance, as our model is defined using proportional abundances instead of the usual number of individuals. To solve this problem we show (see Supplementary Information) that an approximation for the abundance function, defined as the average number of species containing *n* individuals, *n* ∈ {1, …, *J*}, or SAD is:

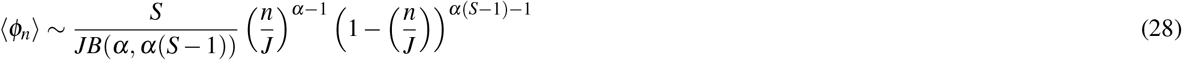

As shown in Table S1 (Supplementary Information) the approximation to the SAD derived from our model is as good as previous ones.

Finally, it is worth reiterating that the form of the stationary distribution *ρ*_∞_ is dependent upon the transitions probabilities characterizing the birth and death process and that the Beta distribution is valid only for the transitions specified by^28^ but other are possible^23,31^. It remains to be seen what other stationary distributions can be found and if these are compatible with observed SADs. This will certainly improve our understanding of the causes underlying the distribution of abundance in ecological systems.

## Acknowledgements

We acknowledge support from projects FONDECYT 1161023, PIA-CONICYT-ACT1112 “Stochastic Analysis Research Network” and VRI-PUC Program on Biostochastics. PAM also acknowledges support from projects ICM-MINECOM, P05-002 IEB, Programa de Financiamiento Basal, CONICYT PFB-23, PIA-CONICYT-Chile, Anillo SOC-1405, and the Santa Fe Institute for providing a stimulating environment while finishing writing the final version of the manuscript. We thank Sergio Rojas who wrote the numerical simulation programs for us and Aurora Gaxiola for comments on the final manuscript and help with the statistical analysis. The python codes used to do Figure 4 are available upon request from the corresponding author.

## Author contributions statement

PAM, GE, AG and RR conceived the study, carried out the mathematical analysis and wrote the manuscript. SRA carried out the statistical fits and data analysis and helped writing the manuscript. All authors reviewed the manuscript.

## References

1. Anderson, W. W. & King, C. E. Age-specific selection. Proceedings of the National Academy of Sciences 66, 780–786(1970).

2. Charlesworth, B. Selection in populations with overlapping generations. i. the use of malthusian parameters in population genetics. Theoretical Population Biology 1, 352–370(1970).

3. Antonovics, J. The input from population genetics:” the new ecological genetics”. Systematic Botany 233–245 (1976).

4. Agrawal, A. A. Community genetics: new insights into community ecology by integrating population genetics. Ecology 84, 543–544(2003).

5. Vellend, M. Species diversity and genetic diversity: parallel processes and correlated patterns. The American Naturalist 166, 199–215(2005).

6. Vellend, M. & Geber, M. A. Connections between species diversity and genetic diversity. Ecology Letters 8, 767–781(2005).

7. Hubbell, S. The Unified Theory of Biodiversity and Biogeography (Princeton University Press, 2001).

8. Volkov, I., Banavar, J. R., Hubbell, S. P. & Maritan, A. Neutral theory and relative species abundance in ecology. Nature 424, 1035–1037(2003).

9. Connolly, S. R., Hughes, T. P. & Bellwood, D. R. A unified model explains commonness and rarity on coral reefs. Ecology Letters 20, 477–486(2017).

10. Haegeman, B. & Etienne, R. S. A general sampling formula for community structure data. Methods in Ecology and Evolution. doi: 10.1111/2041-210X.12807.

11. Etienne, R. S. & Alonso, D. Neutral community theory: how stochasticity and dispersal-limitation can explain species coexistence. Journal of Statistical Physics 128, 485–510(2007).

12. Rosindell, J., Hubbell, S. P., He, F., Harmon, L. J. & Etienne, R. S. The case for ecological neutral theory. Trends in Ecology and Evolution 27, 203–208(2012).

13. Marquet, P. A. et al. On theory in ecology. BioScience 64, 701–710(2014).

14. Engen, S., Solbu, E. B. & Sæther, B.-E. Neutral or non-neutral communities: temporal dynamics provide the answer. Oikos 126, 318–331(2017).

15. Hu, X.-S., He, F. & Hubbell, S. P. Neutral theory in macroecology and population genetics. Oikos 113, 548–556(2006).

16. Leigh, E. G. Neutral theory: a historical perspective. Journal of Evolutionary Biology 20, 2075–2091(2007).

17. Watterson, G. A. Models for the logarithmic species abundance distributions. Theoretical Population Biology 6, 217–250(1974).

18. Blythe, R. A. & McKane, A. J. Stochastic models of evolution in genetics, ecology and linguistics. Journal of Statistical Mechanics: Theory and Experiment 2007, P07018 (2007).

19. de Vladar, H. P. & Barton, N. H. The contribution of statistical physics to evolutionary biology. Trends in Ecology and Evolution 26, 424–432 (2011).

20. Feller, W. Diffusion processes in genetics. In Second Symposium on Probability and Statistics (University of California Press Berkeley, Calif., 1951).

21. Vallade, M. & Houchmandzadeh, B. Analytical solution of a neutral model of biodiversity. Physical Review E 68, 061902 (2003).

22. McKane, A. J., Alonso, D. & Solé, R. V. Analytic solution of hubbell’s model of local community dynamics. Theoretical Population Biology 65, 67–73 (2004).

23. Etienne, R. S., Alonso, D. & McKane, A. J. The zero-sum assumption in neutral biodiversity theory. Journal of Theoretical Biology 248, 522–536 (2007).

24. Allen, A. P. & Savage, V. M. Setting the absolute tempo of biodiversity dynamics. Ecology letters 10, 637–646 (2007).

25. Wright, S. Evolution in mendelian populations. Genetics 16, 97–159 (1931).

26. Wright, S. The distribution of gene frequencies in populations. Proceedings of the National Academy of Sciences 23, 307–320 (1937).

27. Rebolledo, R. La méthode des martingales appliquée à l’étude de la convergence en loi de processus. Mémoires de la Societe Mathematique de France 62, 1–129 (1979).

28. McKane, A., Alonso, D. & Solé, R. V. Mean-field stochastic theory for species-rich assembled communities. Physical Review E 62, 8466 (2000).

29. Fisher, R. A., Corbet, A. S. & Williams, C. B. The relation between the number of species and the number of individuals in a random sample of an animal population. The Journal of Animal Ecology 42–58 (1943).

30. Terborgh, J., Robinson, S. K., Parker, T. A., Munn, C. A. & Pierpont, N. Structure and organization of an amazonian forest bird community. Ecological Monographs 60, 213–238 (1990).

31. Volkov, I., Banavar, J. R., He, F., Hubbell, S. P. & Maritan, A. Density dependence explains tree species abundance and diversity in tropical forests. Nature 438, 658–661 (2005).

32. Latimer, A. M., Silander, J. A. & Cowling, R. M. Neutral ecological theory reveals isolation and rapid speciation in a biodiversity hot spot. Science 309, 1722–1725 (2005).

33. Dornelas, M., Connolly, S. R. & Hughes, T. P. Coral reef diversity refutes the neutral theory of biodiversity. Nature 440, 80–82 (2006).

34. Etienne, R. S., Latimer, A. M., Silander, J. A. & Cowling, R. M. Comment on “neutral ecological theory reveals isolation and rapid speciation in a biodiversity hot spot”. Science 311, 610b–610b (2006).

35. Wright, S. Evolution and the Genetics of Populations, volume 2: The theory of gene frequencies (University of Chicago Press, 1969).

36. Engen, S. & Lande, R. Population dynamic models generating the lognormal species abundance distribution. Mathematical Biosciences 132, 169–183 (1996).

37. Pigolotti, S., Flammini, A. & Maritan, A. Stochastic model for the species abundance problem in an ecological community. Physical Review E 70, 011916 (2004).

38. Azaele, S., Pigolotti, S., Banavar, J. R. & Maritan, A. Dynamical evolution of ecosystems. Nature 444, 926–928 (2006).

39. Kolmogorov, A. N. Deviations from hardy’s formula in partial isolation. In Comptes Rendus de l’Academie des Sciences de l’URSS Nouvelle Serie, vol. 3, 129–132 (1935).

40. Leigh, E. G. Ecological role of volterra’s equations. In Gerstenhaber, M. (ed.) Some Mathematical Problems in Biology, 1–61 (American Mathematical Society, Providence, Rhode Island, USA, 1968).

41. Dennis, B. & Patil, G. The gamma distribution and weighted multimodal gamma distributions as models of population abundance. Mathematical Biosciences 68, 187–212(1984).

42. Engen, S. & Lande, R. Population dynamic models generating species abundance distributions of the gamma type. Journal of Theoretical Biology 178, 325–331 (1996).

43. Kempton, R. A generalized form of fisher’s logarithmic series. Biometrika 29–38 (1975).

44. Boyer, J. F. The effects of prior environments on tribolium castaneum. The Journal of Animal Ecology 865–874 (1976).

45. Losos, J. B. & Adler, F. R. Stumped by trees? a generalized null model for patterns of organismal diversity. The American Naturalist 145, 329–342(1995).

46. Ogle, K. et al. Quantifying ecological memory in plant and ecosystem processes. Ecology letters 18, 221–235(2015).

47. Ewens, W. J. Mathematical Population Genetics 1: Theoretical introduction, vol. 27 (Springer-Verlag, New York, 2004).

48. Kimura, M. Diffusion models in population genetics. Journal of Applied Probability 1, 177–232(1964).

49. Lau, H.-S. & Hing-Ling Lau, A. Effective procedures for estimating beta distribution’s parameters and their confidence intervals. Journal of Statistical Computation and Simulation 38, 139–150(1991).

50. Dacunha-Castelle, D. & Florens-Zmirou, D. Estimation of the coefficients of a diffusion from discrete observations. Stochastics: An International Journal of Probability and Stochastic Processes 19, 263–284(1986).

51. Rebolledo, R. Open quantum systems and classical trajectories. In Rebolledo R, Z. J., Rezende J (ed.) Stochastic Analysis and Mathematical Physics: The mathematical legacy of RP Feynman, 141–164 (World Scientific Publishing, Singapore, 2004).

